# On the Apportionment of Archaic Human Diversity

**DOI:** 10.1101/2021.07.15.452563

**Authors:** Kelsey E. Witt, Fernando Villanea, Elle Loughran, Emilia Huerta-Sanchez

## Abstract

The apportionment of human genetic diversity within and between populations has been measured to understand human relatedness and demographic history. Likewise, the distribution of archaic ancestry in modern populations can be leveraged to better understand the interaction between our species and its archaic relatives, and the impact of natural selection on archaic segments of the human genome. Resolving these interactions can be difficult, as archaic variants in modern populations have also been shaped by genetic drift, bottlenecks, and gene flow. Here, we investigate the apportionment of archaic variation in Eurasian populations. We find that archaic genome coverage at the individual- and population-level present unique patterns in modern human population: South Asians have an elevated count of population-unique archaic SNPs, and Europeans and East Asians have a higher degree of archaic SNP sharing, indicating that population demography and archaic admixture events had distinct effects in these populations. We confirm previous observations that East Asians have more Neanderthal ancestry than Europeans at an individual level, but surprisingly Europeans have more Neandertal ancestry at a population level. In comparing these results to our simulated models, we conclude that these patterns likely reflect a complex series of interactions between modern humans and archaic populations.

## Introduction

In *The Apportionment of Human Diversity*, R. C. Lewontin endeavored to address the partition of genetic diversity within and between human populations. This work was fundamental at a time when human population structure was the domain of morphology — a field long influenced by colonialism and Eurocentrism. Lewontin’s work paved the way for our current understanding of human population structure: a continuous gradient of diversity that was influenced by human migrations originating in the African continent [1], as populations that are geographically further from Africa have fewer variable sites and lower heterozygosity [2–4]. Additionally, more recent periods of population replacement [5,6] or gene flow [7], isolation, and selective pressures [8,9] have further shaped the genomes of modern populations.

Since Lewontin’s groundbreaking work, an additional component of human genetic diversity has been discovered and highlighted in recent decades: modern human populations carry a legacy of admixture with archaic human populations, including Neanderthals and Denisovans. Neanderthal ancestry has been detected in human populations in Eurasia, Oceania, and the Americas, as well as North Africans [10–12], while Denisovan ancestry has been found primarily in Asia, the Americas, and Oceania [11,13,14]. Further archaic ancestry from unknown sources has even been identified in African populations [15–17]. Levels of archaic ancestry as a whole (including Neanderthal and Denisovan introgression, as well as other archaic humans in the case of Africans) vary between ∼1% in African populations [15,16] and ∼2% in Eurasians [11,13,14], with populations in Oceania harboring the largest amount at ∼6% [14,36]. The surviving archaic segments in modern human genomes are likely not the product of a single admixture event, but instead reflect a complex history of multiple points of contact between humans and several archaic populations [16,18–20]. Interbreeding with archaic humans introduced new genetic variation into modern humans, and these archaic variants were shaped by demographic and selective forces. Positive [21–23] and negative [11,24–26] selection have shaped the frequency of some archaic genome segments, but genetic drift amplified by demographic processes — population contractions and expansions — along with admixture between modern human lineages are largely responsible for the current distribution of archaic variation in modern populations [27]. Gene flow from populations with population-unique archaic alleles can introduce new archaic variants to a population, or gene flow from populations without archaic admixture can decrease the amount of archaic ancestry in a population [28,29].

One key observation related to the apportionment of archaic ancestry is that despite most Neanderthal archeological sites being situated in western Eurasia, East Asian individuals exhibit higher Neanderthal ancestry than modern Europeans [10,11,30,31]. Some studies have suggested that differences in demography between East Asians and Europeans are sufficient to explain the elevated Neanderthal ancestry in East Asians [11,30,32]. Other studies have found that these factors explain some but not all of the difference [19,25], suggesting instead that additional Neanderthal admixture events provide a better explanation for the observed patterns in modern populations [14,19,33]. Interestingly, a study that examined the genetic differentiation between archaic ancestry segments in different populations recovered signals from two distinct Denisovan populations but only one Neanderthal population [18]. This suggests that if Neanderthal admixture did occur more than once, it was from the same population or multiple closely-related ones. Europeans, however, have a complex history, and this may have affected levels of Neanderthal ancestry. The earliest Europeans, who encountered European Neanderthals, are more closely related to East Asians [34], and were replaced by later migrants after all Neanderthals had become extinct [35]. Europeans further received gene flow from other Eurasian populations [28,37], and maintained long-term gene flow with African populations [15,38]. Because of the complexity of Eurasian demographic history, none of these studies have found one single cause for the differences in Neanderthal ancestry between these populations. Instead, the evidence points toward a more complex interaction of population demography, natural selection, and possibly multiple admixture events.

Several previous studies have inferred and quantified levels of archaic ancestry in modern human populations, and in this study we wanted to look more closely at the patterns of archaic variation in each population to gain insight into how this variation has evolved in modern humans. Specifically, we compute the apportionment and frequency of archaic variation in human populations and we quantify levels of shared and non-shared archaic variation between modern populations. We find that, similarly to non-archaic variants, the majority of Neanderthal variation is shared between populations. Denisovan variation, however, is mostly unique to specific populations. Archaic variation in a population has also been impacted by its demographic history; for example, observable population structure from archaic variants mirrors that of non-archaic variants. We also quantify the level of archaic variation as a function of sample size, and we find that more of the Neanderthal genome can be recovered from a sample of South Asian individuals than a sample (of equal size) of Europeans or East Asians. In comparing Europeans with East Asians, we confirm that East Asian individuals harbor a larger amount of Neanderthal ancestry than European individuals, as previously reported, but more of the Neanderthal genome is recovered from a sample of Europeans than an equal size sample of East Asians individuals. We use simulations to explore demographic models of archaic introgression and assess which model is most consistent with the patterns observed in the empirical data. Examining the apportionment of archaic ancestry at the population level will improve our understanding of how differing demographic histories have impacted the distribution and number of archaic alleles in modern human populations.

## Methods

### Archaic genome coverage

To study patterns of archaic variation in modern human populations, we examined the quantity and the frequency of archaic introgressed variants. Using the autosomes, we measured the amount of archaic ancestry within a single individual as well as in a set of multiple individuals. We call this measure “archaic genome coverage”, and use it to investigate how sample size impacts the quantity of an archaic genome recovered. We computed this quantity by using the number of single nucleotide polymorphisms (SNPs) that we identify as “archaic”, which is defined in the next section. Figure 1 illustrates our concept of archaic genome coverage at the individual and population level. Here, we show the archaic genome coverage in a genome region for two populations (A and B), each containing four individuals. The individual-level archaic coverage is simply the number of archaic SNPs identified across the genome. For the genome region in our example, the genome coverage for individuals in population A ranges from 3-4 and the genome coverage for individuals in population B ranges from 1-2. To take the genome coverage of a larger number of individuals, we look for all sites where at least one individual in the sample has an archaic SNP. Therefore, population A has archaic genome coverage of 5 and population B has archaic genome coverage of 6. Our example also illustrates how population- and individual-level genome coverage can vary between populations.

**Figure 1:**
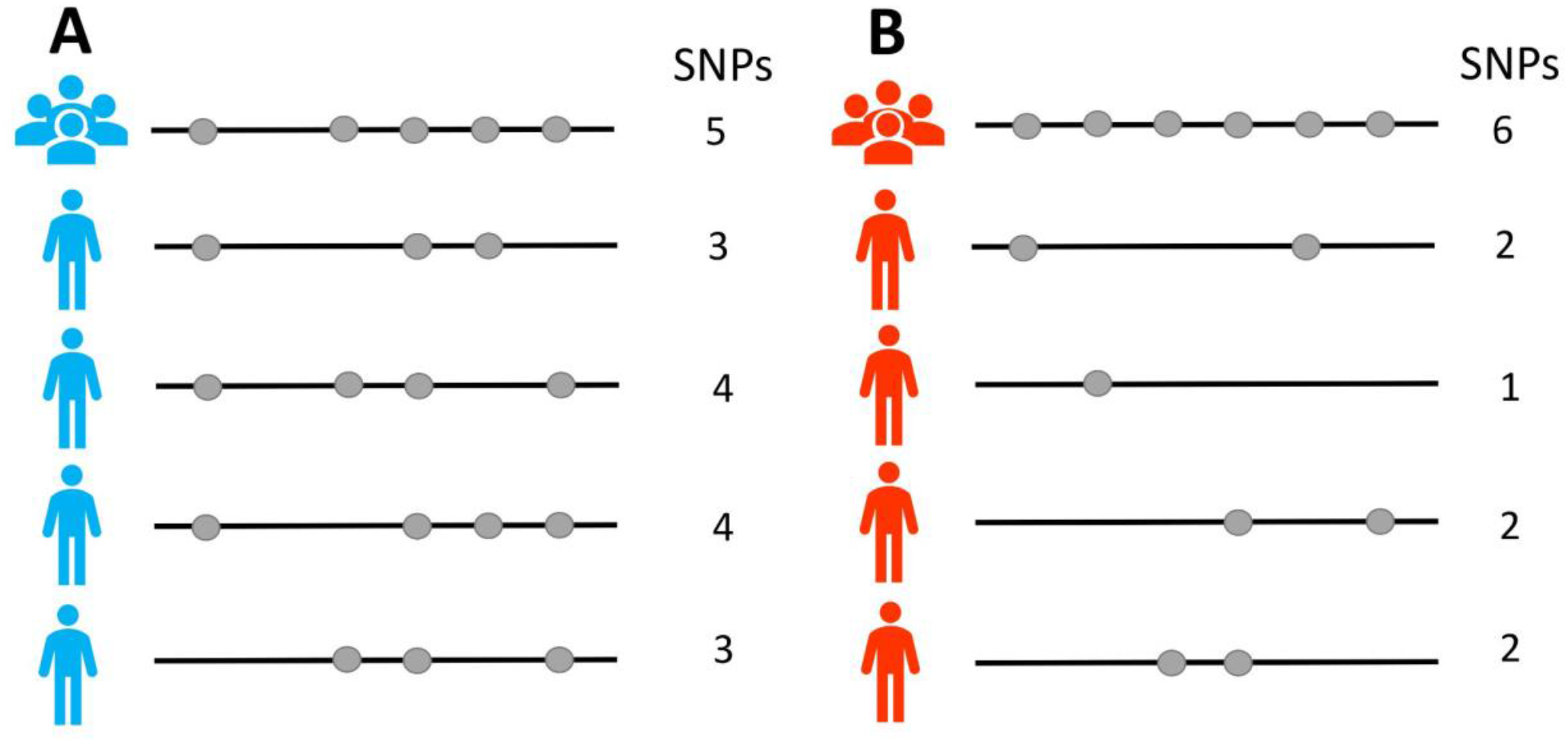
An illustration of population- and individual-level archaic genome coverage. Here we show the archaic SNPs (gray circles) present in a genomic region (the black line) for two populations, A and B. Each population contains four individuals, and their genome coverage is shown next to each individual along with the total number of SNPs they have. For the population-level coverage, each archaic SNP that is found in any individual in the population counts towards the total, so population-level coverage is the sum of archaic SNPs found across all individuals in that population (the top line in A and B).

Population B has higher genome coverage at the population level, but lower genome coverage at the individual level, suggesting that there is more archaic allele sharing between individuals within population A. Note that this is similar to counting the number of segregating sites (*S*), except we are conditioning on the mutations also being shared with Neanderthals or Denisovans. This analysis can similarly be applied to ancestry tracts instead of SNPs, provided that the ancestry tracts are identified at the individual level (Supplemental Figure 1)

### Identifying archaic SNPs and calculating genome coverage

We compared the 1000 Genomes (Phase III) populations [39] to the Altai [40], Vindija [41], and Chagyrskaya Neanderthal [42], and the Denisovan [36] high-coverage genomes. Archaic genotypes were filtered with a minimum genotype quality score of 40. SNPs were considered to be “Non-African” if two conditions were true: 1) the allele at that SNP had a frequency less than 0.01 across all African 1000 Genome populations and 2) the allele had a frequency greater than 0.01 in at least one non-African population. These two conditions were set to identify sites with mutations that most likely arose outside of Africa. In addition, if the allele at that SNP was also found in at least one of the sequenced archaic genomes, then we call it an archaic SNP, to represent sites with mutations that were likely introgressed from archaic humans. The Non-African SNPs that were not archaic were defined as *Modern Non-African* SNPs, which have the same allele frequency requirements as the archaic datasets but are not shared with archaic individuals. We excluded two populations — ACB (African Caribbeans in Barbados) and ASW (African Ancestry in Southwest US) — from our analyses because they contain a high proportion of African ancestry, so we expect them to have low levels of Neanderthal or Denisovan ancestry. We considered three sets of archaic SNPs: *All-Archaic* (found in any of the archaic genomes), *Denisovan-Unique* (found in the Denisovan but no Neanderthals), and *Neanderthal-Unique* (found in the Altai, Chagyrskaya, or Vindija Neanderthals but not Denisovans). We further examined archaic allele sharing between populations and geographic regions (Europe, East Asia, South Asia, and the Americas), to count the number of archaic alleles in the All-Archaic, Neanderthal-Unique, and Denisovan-Unique sets that were unique to a population or region or were found worldwide.

We counted all SNPs with Non-African alleles present in each population and partitioned them as modern or archaic (see definitions in previous paragraph). For each of these modern and archaic alleles, we calculated the allele frequency and classified them as “rare” (.01<f<0.2) or “common” (f>= 0.2) to examine patterns in allele frequency distribution. We further partitioned archaic variants into Neanderthal-Unique, Denisovan-Unique, and All-Archaic to estimate the contribution of each archaic hominin to the archaic ancestry present in each population. We calculated the archaic genome coverage per individual by summing up the total number of archaic SNPs in each individual’s genome (Figure 2A, 2C, 2E). We computed the archaic genome coverage in samples of randomly-selected individuals from each population of varying size (n=1,10, 25, 50, 75, 100, 125, 150, see Figure 2B, 2D, 2F).

**Figure 2:**
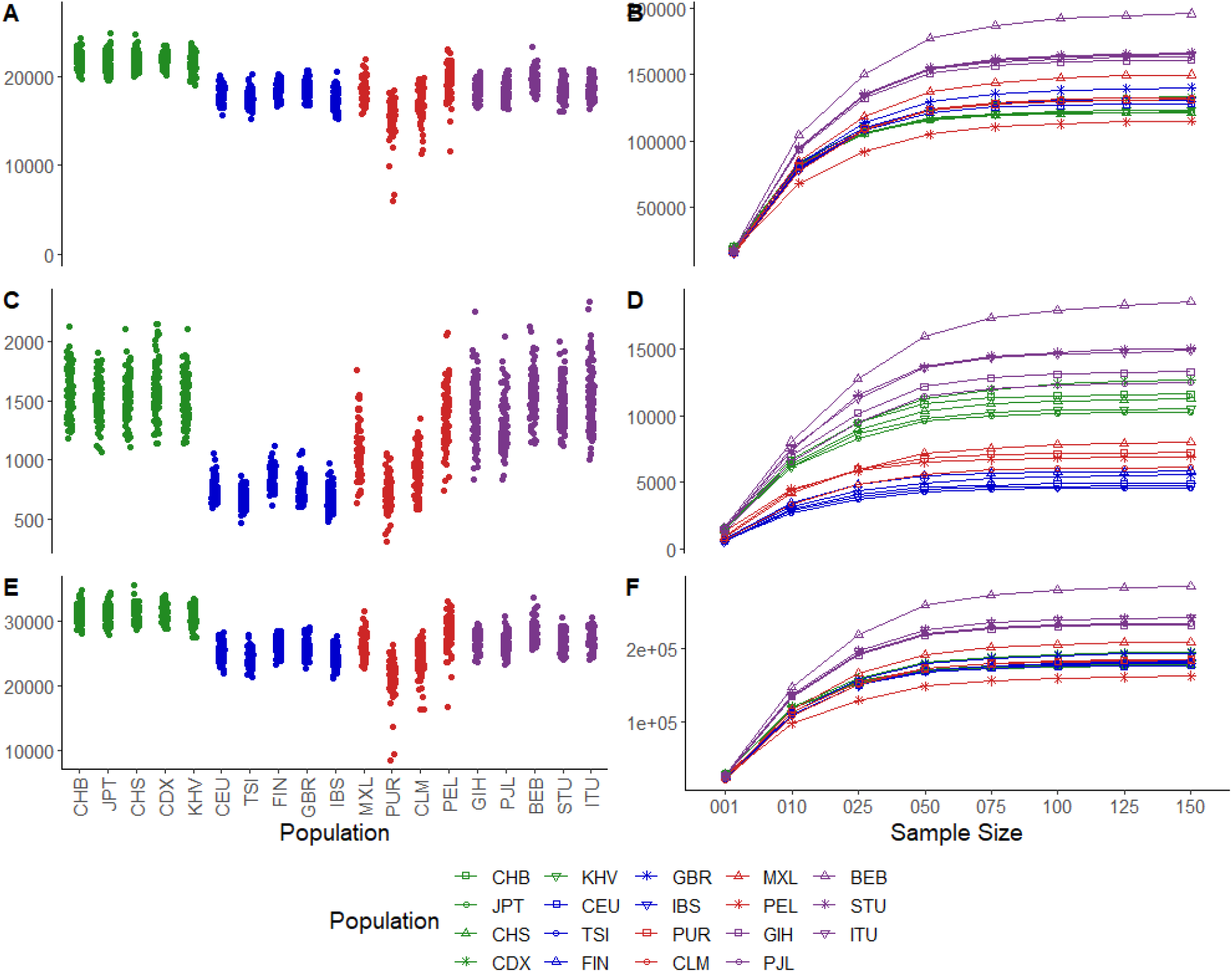
Individual archaic genome coverage (GC) counts for Neanderthal-Unique (A), Denisovan-Unique (C), and All-Archaic (E) SNPs in the 1000 Genome Populations in East Asia (green), Europe (blue), the Americas (red), and South Asia (purple), and mean values for genome coverage of each population at varying sample sizes (n=1,10, 25, 50, 75, 100, 125, 150) for Neanderthal-Unique (B), Denisovan-Unique (D), and All-Archaic (F) SNPs. The genome coverage values for n=1 on plots B, D, and F are the median values for each population in plots A, C, and E. Populations are color-coded by region and abbreviations follow standard conventions established for the 1000 Genomes Project data.

For some analyses, namely the case of Neanderthal introgression into Europeans and East Asians, we also computed the archaic genome coverage using the introgressed tract lengths inferred in other studies [11,18,31]. For the studies that included SNP data [18,31], we counted the archaic SNPs as identified by each of the studies that were present in the 1000 Genomes CEU, CHB, and CHS populations. For the Sankararaman et al. (2014) dataset that used introgressed tracts rather than SNPs [11], we used the introgressed haplotypes for CHB, CHS, JPT, IBS, TSI, CEU, FIN and GBR 1000 Genome Project individuals, excluding X chromosome haplotypes. To compare Neanderthal genome coverage across European and East Asian super populations, we sampled haploid individuals and merged introgressed tracts between haploids in each sample using the merge function in BEDtools version 2.26 [43] to find the total length of Neanderthal genome recovered. We took one hundred replicates of each of nine sample sizes (1, 5, 10, 25, 50, 75, 100, 125, 150 haploid individuals) from each superpopulation to calculate the ratio of European to East Asian Neanderthal genome coverage. We also compared homozygosity of Neanderthal introgressed tracts between European and East Asian individuals by pairing haplotypes as identified in [11] into their diploid individuals and identified intersections between tracts on each allele for each diploid using the intersect function in BEDtools version 2.26. We considered a tract homozygous if there was a tract on its paired allele that reciprocally overlapped it by at least a threshold percentage (40, 50, 60, 70, 75, 80, 90 or 95%, see supplemental Figure 8).

### Principal component analysis

To determine if archaic variants in humans can be used to reproduce known patterns of human population structure, we used principal component analysis (PCA). We used the archaic SNPs (see definition in section titled “Identifying archaic SNPs and calculating genome coverage”) with a minimum frequency of 0.05 in at least one non-African population for the PCA (n=∼250,000). We also selected an equal number of randomly-selected Non-African Modern SNPs (which had a frequency in Africans of less than 0.01 and a minimum frequency of 0.05 in a non-African population) to serve as a non-archaic comparison. PCAs were constructed for all three sets of archaic SNPs (All-Archaic, Neanderthal-Unique, and Denisovan-Unique, see Figure 3A-C) and the randomized Modern SNP subset (Supp. Figure 2) using Eigenstrat version 6.0 [44,45]. The resulting PCAs were plotted in R version 4.0.2 using ggplot2 [46,47].

**Figure 3:**
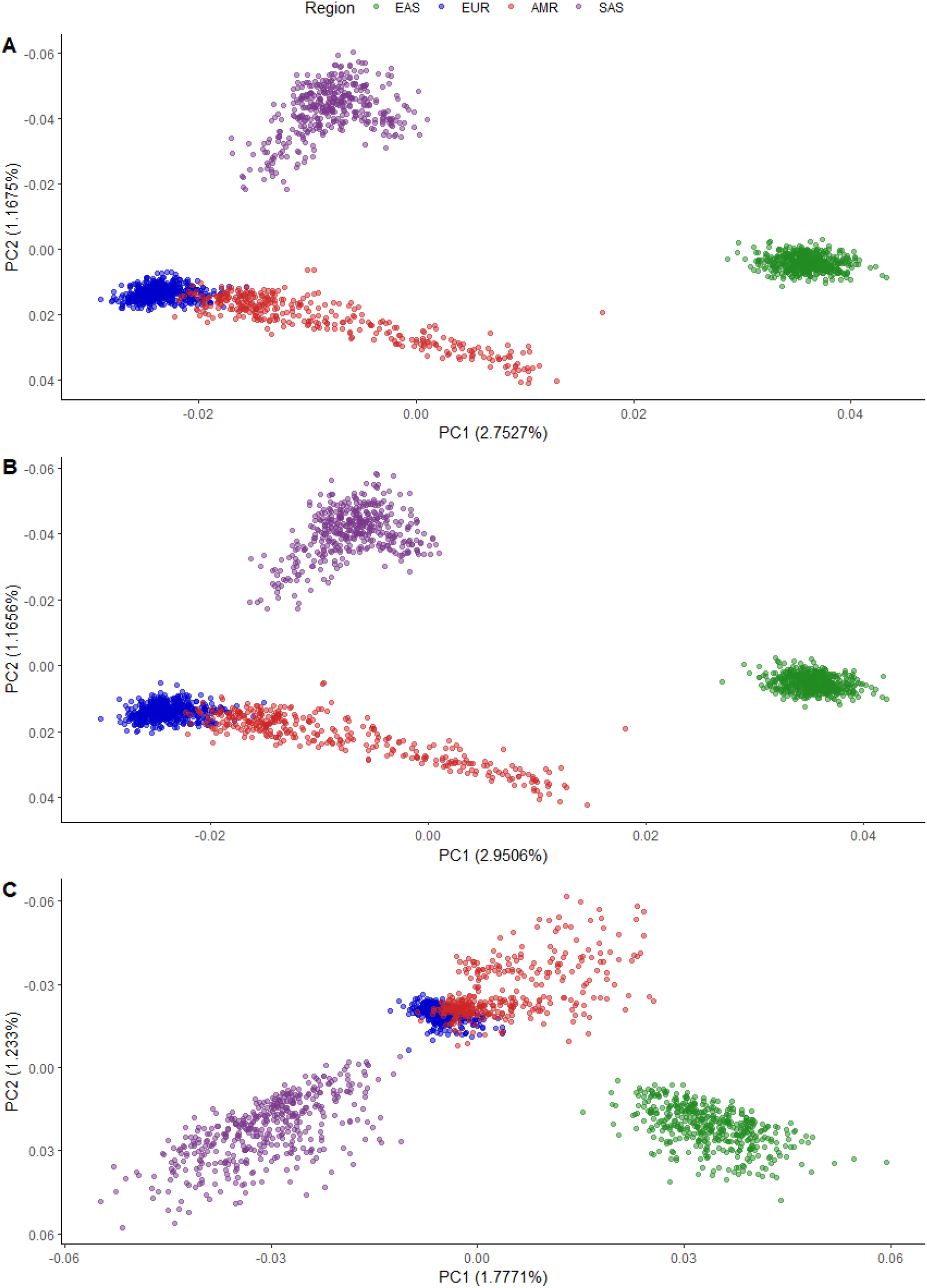
A PCA of 1000 Genomes populations, using archaic SNPs with a frequency of at least 5% in one non-African population for A) All-Archaic SNPs, B) Neanderthal-Unique SNPs, and C) Denisovan-Unique SNPs. Individuals are color-coded by their super-population: EAS (East Asians), EUR (Europeans), AMR (Americans), SAS (South Asians).

### Simulations in MSprime

To investigate a set of proposed demographic scenarios that may be responsible for the observed relationship between the amount of Neanderthal ancestry recovered as a function of sample size (Figure 1, Figure 4), we used msprime version 0.7.2 [48], to simulate archaic introgression into modern Europeans and East Asians. Specifically, we wanted to test whether one or two introgression events could lead to this pattern. For the simulations we fixed demographic parameters, including effective population sizes and divergence times, based on the Gravel model [49,50], extended to accommodate archaic introgression based on Villanea and Schraiber ([14,19,33]). All fixed parameters are listed in Supp. Figure 3. In order to explore how various levels of admixture with archaic populations impact the amount of archaic variation recovered (archaic genome coverage) in modern populations, we tested two admixture parameters: a “first pulse” of Neanderthal gene flow into the ancestor of Europeans and East Asians (where admixture proportions are: 1%, 1.5%, 2%, 2.5%, 3 or 4%) and a “second pulse” of Neanderthal gene flow into East Asians (where admixture proportions are: 0%, 0.1%, 0.2%, 0.5%, 0.8%, 1%) following the East Asian-European split (Supp. Figure 3).

**Figure 4:**
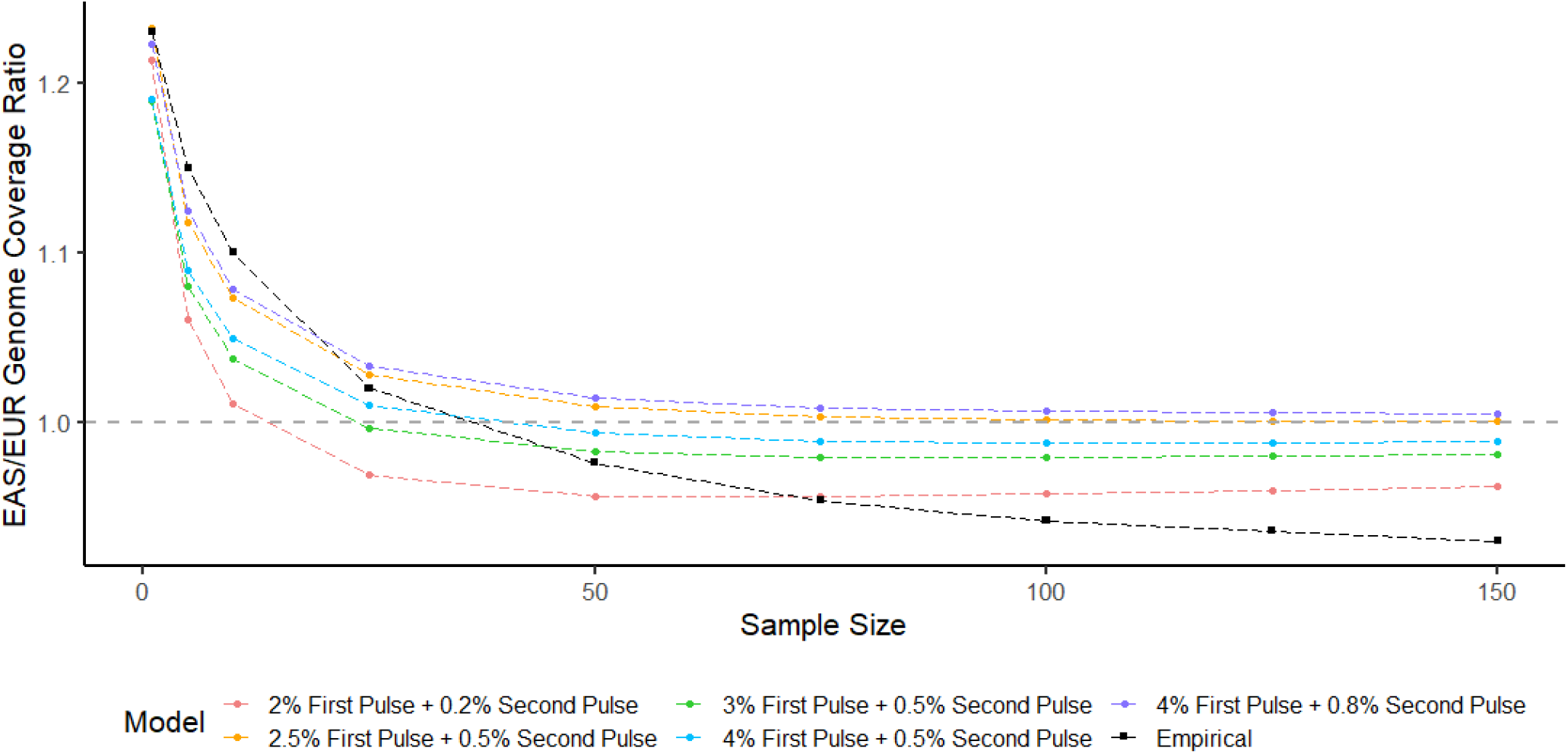
Comparing archaic genome coverage in East Asians (EAS) and Europeans (EUR) across simulated and empirical datasets. The x axis is the number of individuals sampled to calculate genome coverage, and the y axis is the genome coverage found in EAS divided by the genome coverage found in EUR. The dashed horizontal line denotes where the genome coverage would be equal across both populations. The empirical mean values (from 100 sampled replicates) are in black, and the mean values (from 100 sampled replicates each of 200 simulated datasets) of the five models with the lowest mean squared error relative to the empirical data are shown in different colors. For all models, the “First” pulse represents gene flow from Neanderthals into the ancestor of East Asians and Europeans, while the “Second” pulse represents archaic gene flow into East Asians specifically. The full list of models, their coverage ratio values, and mean squared error is available in Supplemental Table 1.

For each replicate, we simulated chromosomes of 85 European individuals (170 chromosomes), and 198 East Asian individuals (396 chromosomes), matching the sampling available from the 1000 Genomes Project panel for the CEU and CHB+CHS populations (the latter two populations were combined because of their high genetic similarity). We simulated a 100Mb chromosome using a mutation rate of 1.5e-8 bp/gen and a recombination rate of 1e-8 bp/gen. Using the tree sequences output by msprime, we identified introgressed segments in the sampled chromosomes by asking which of the sampled chromosomes coalesced with the archaic lineage more recently than the human-archaic population split time. For each simulation replicate we computed the amount of Neanderthal recovered in the simulated Europeans and East Asians populations as a function of the sampled chromosomes, and took the ratio of East Asian archaic genome coverage to European archaic genome coverage (EAS/EUR). Each combination of admixture parameters was simulated with 200 replicates. For each replicate, we resampled genomes 100 times for each sample size. For example, for a sample of size 1, we randomly sampled one European chromosome and one East Asian chromosome, took the ratio, and we did that 100 times and computed the mean across all replicates.

For our empirical data comparison, we calculated the ratio of East Asian to European archaic genome coverage using the Neanderthal-Unique SNP set across various sample sizes (n=1, 10, 25, 50, 75, 100, 125, 150), resampling the data 100 times for each sample size to create a distribution. We also compared our results to that of three previously-published datasets: the archaic SNPs identified using the method Sprime in Browning et al. [18], the archaic SNPs identified using the program DICAL-ADMIX in [31], and the archaic introgressed tracts identified using a conditional random field method in Sankararaman et al. [11]. A comparison of the empirical archaic genome coverage is shown in Supp. Figure 4. For each dataset, we calculated the ratio of East Asian to European genome coverage at the sample sizes mentioned above (using SNPs or tract lengths depending on the data), resampling 100 times for each size. Because our simulated data produced tract lengths, we chose to compare our simulated data to the inferred introgressed maps from Sankararaman et al. (2014). We calculated the ratio of East Asian to European archaic genome coverage across sample sizes for each of the simulated datasets. We calculated mean squared error to test the fit of each model to the empirical data:

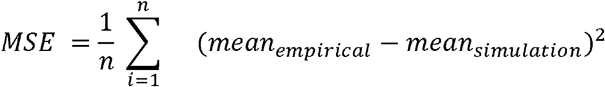

where n= the number of simulation replicates for each model.

## Results

### Patterns of archaic variation in modern human populations

We examined the apportionment and frequency of archaic alleles across modern human populations, and asked if patterns of archaic genome coverage change as a function of the sample size; when we are examining individual- or population-level samples (Figure 1). As a first step, we confirmed that archaic genome diversity was shaped by the same demographic forces as the rest of the genome by applying a principal component analysis (PCA) to the set of All-Archaic sites (see Methods) with archaic alleles at >5% frequency. We find that archaic alleles perform as well as a size-matched random sample of non-archaic variants by recapitulating similar levels of population structure as that obtained from non-archaic sites (Figure 3A). Archaic alleles carry sufficient information to visually distinguish between East Asian, South Asian and European populations. The first principal component visually separates East Asians, South Asians and Europeans, while the second principal component differentiates the admixed American, European and East Asian populations from the South Asian populations. As expected, the first principal component also sorts the admixed American populations based on their proportion of European ancestry, so that individuals with higher European ancestry cluster more closely with Europeans (Supplemental Figure 5). Neanderthal-Unique sites show a similar pattern to that of All-Archaic sites (Figure 3B), while Denisovan-Unique sites show less distinction between South Asians and Europeans (Figure 3C).

**Figure 5:**
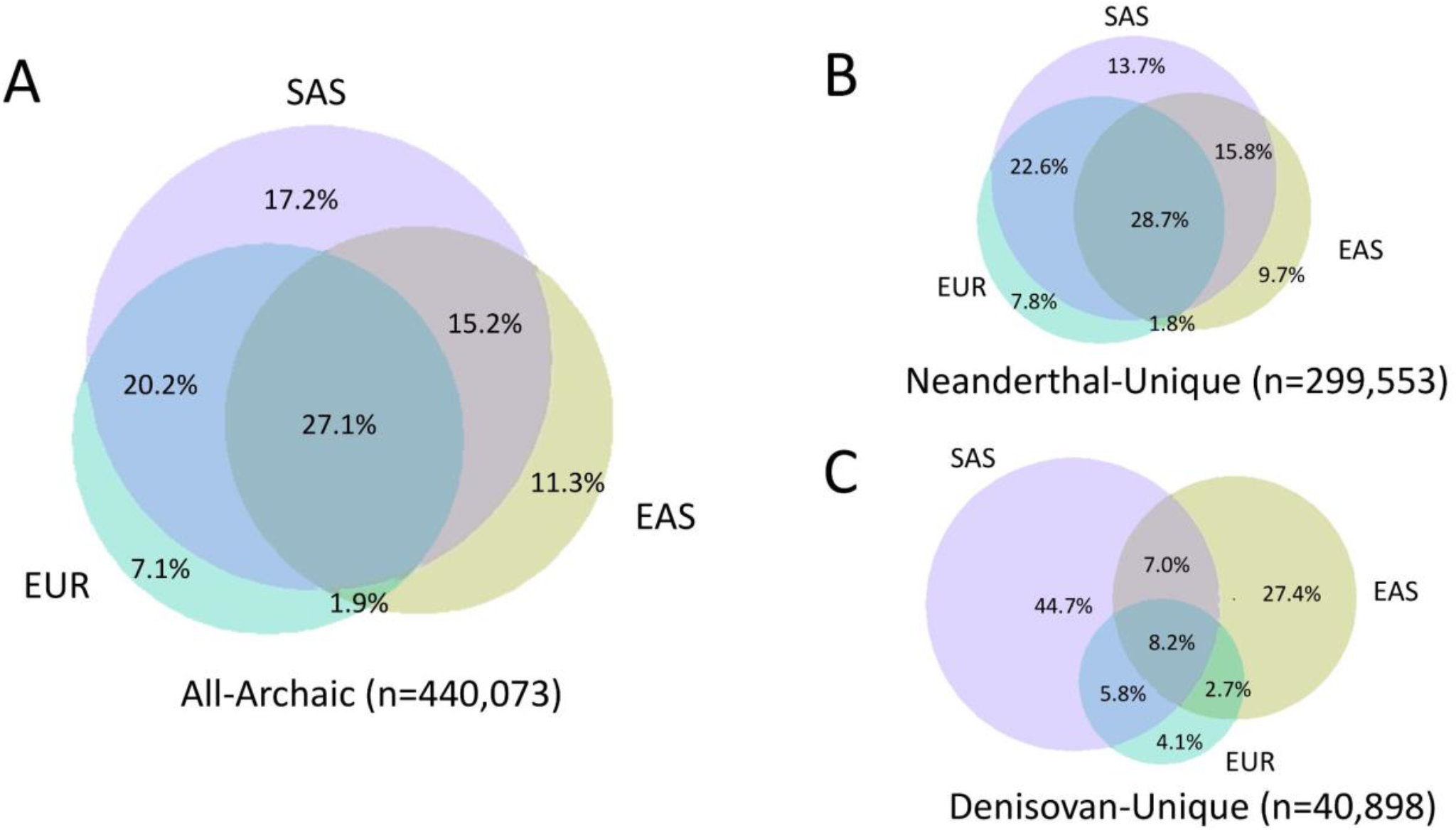
A Venn Diagram showing archaic allele sharing between geographic regions in Eurasia: Europeans (EUR), East Asians (EAS), and South Asians (SAS) for A) All-Archaic alleles, B), Neanderthal-Unique alleles, and C) Denisovan-Unique alleles. The total number of SNPs in each dataset is included below each plot, and the percentages refer to the percentage of SNPs shared by the populations in overlapping circles.

Despite the regional differences as observed in the PCA, there is more variation that is shared between populations and regions than is population- or region-unique (Table 1) — consistent with Lewontin’s [51] original observations, as well as more recent research <u>(2,52)</u>. For example, when we examine archaic allele sharing between Eurasian populations (East Asians, South Asians and Europeans), we find that ∼64% of all archaic alleles are shared by at least two populations (Figure 5A). Archaic variants present in only a single Eurasian population make up 35.6% of archaic variants, with 17.2% of them found in South Asians, 11.3% found in East Asians and 7.1% found in Europeans. These numbers show that South Asians have the largest number of unique archaic alleles relative to other Eurasian populations (17.2%). If we examine only the Neanderthal-Unique alleles (Figure 5B), the trends are similar to those observed for all archaic alleles (Figure 5B). Notably, the Denisovan-unique alleles show a different pattern, where only 23.7% of all Denisovan-unique SNPs are found in at least two populations, and a large proportion (76.2%) of Denisovan-unique variation is private to South Asian or East Asian populations (44.7% and 27.4% respectively, see Figure 5C). This may be a consequence of contributions from distinct Denisovan populations into East and South Asians.

**Table 1:** Counts of all archaic alleles, as well as the number of archaic alleles shared across all super-populations (East Asians [EAS], Europeans [EUR], Americas [AMR], and South Asians [SAS],n=448512) and all populations, as well as the count of archaic alleles that are found only in a single super-population or population. In this case “unique” signifies that the variant is only present in a single super-population or population.

| Total |  | Super-populations |  | Populations |  |
| --- | --- | --- | --- | --- | --- |
| All populations | 448512 | All super-populations | 107914 | All populations | 38604 |
|  |  | EAS-unique | 40955 | CHB-unique | 1537 |
|  |  |  |  | CDX-unique | 6156 |
|  |  |  |  | CHS-unique | 1363 |
|  |  |  |  | JPT-unique | 4529 |
|  |  |  |  | KHV-unique | 1844 |
|  |  | EUR-unique | 12098 | CEU-unique | 728 |
|  |  |  |  | FIN-unique | 1702 |
|  |  |  |  | GBR-unique | 2335 |
|  |  |  |  | IBS-unique | 770 |
|  |  |  |  | TSI-unique | 1553 |
|  |  | AMR-unique | 8439 | CLM-unique | 2881 |
|  |  |  |  | MXL-unique | 627 |
|  |  |  |  | PEL-unique | 677 |
|  |  |  |  | PUR-unique | 1688 |
|  |  | SAS-unique | 71461 | BEB-unique | 7830 |
|  |  |  |  | GIH-unique | 2979 |
|  |  |  |  | ITU-unique | 4632 |
|  |  |  |  | PJL-unique | 2522 |
|  |  |  |  | STU-unique | 5181 |

While most of the variation in non-Africans is a subset of what we observed in African populations, non-African populations accrued new mutations since their expansion out of Africa. If we ask, what proportion of these mutations (defined by our “Non-African” set, see Methods) were actually introduced through introgression with archaic humans (i.e mutations are also present in the sequenced archaic individuals), we find that it varies between 7% and 11% depending on the population (see Table 1). As expected, the majority (88-98%) of Non-African alleles, whether looking at the modern (non-archaic) or the archaic set, have rare allele frequencies <20%. The populations with the largest proportions of high frequency (>= 20%) non-archaic or archaic alleles are found in East Asians and Peruvians (6-12% compared to 2-6% for other populations, Tables 1-2). For most populations, the ratio of common to rare alleles is similar regardless of whether the SNPs being considered are archaic or modern (Supplementary Figure 6). The only exception is South Asians, who not only have more archaic variants in the population, but these archaic variants tend to be more rare than in other populations; South Asians have a significantly higher proportion of rare archaic alleles compared to rare modern non-African alleles (Tables 1,2 and Supplemental Figure 6). The pattern of more unique Denisovan variants in South Asians suggests contributions from multiple Denisovan populations and these variants are segregating at lower frequencies perhaps because present-day South Asians are descendants of multiple ancestral populations that harbored contributions from distinct Denisovan populations [53].

**Table 2:** Counts of archaic (Non-African and archaic) alleles and modern (Non-African and non-archaic) alleles as well as the proportions of Neanderthal-unique and Denisovan-unique variants, the percentage of non-African alleles that are archaic, as well as the proportion of rare (<20% frequency) and common alleles (>20% frequency), and the ratio of archaic common percentage to modern common percentage. Populations are referred to using the standard 1000 Genomes convention.

|  | Total Archaic Alleles | Total Modern Alleles | % Neanderthal-Unique | % Denisovan-Unique | % of Non-African archaic Alleles | % Common archaic alleles | % common non-archaic alleles | % Rare archaic alleles | % Rare non-archaic alleles | Archaic/Modern Common Ratio |
| --- | --- | --- | --- | --- | --- | --- | --- | --- | --- | --- |
| CHB | 178685 | 1556889 | 68.46 | 6.53 | 10.71 | 9.48 | 8.31 | 90.52 | 91.69 | 1.14 |
| JPT | 176451 | 1689852 | 68.99 | 5.85 | 9.86 | 9.54 | 7.75 | 90.46 | 92.25 | 1.23 |
| CHS | 177815 | 1556151 | 68.68 | 6.35 | 10.67 | 9.36 | 8.30 | 90.64 | 91.70 | 1.13 |
| CDX | 197348 | 2263587 | 68.40 | 6.56 | 8.40 | 8.42 | 5.84 | 91.58 | 94.16 | 1.44 |
| KHV | 179977 | 1560242 | 68.70 | 5.85 | 10.77 | 9.23 | 8.32 | 90.77 | 91.68 | 1.11 |
| CEU | 177132 | 1911470 | 72.60 | 2.79 | 8.87 | 5.88 | 5.04 | 94.12 | 94.96 | 1.17 |
| TSI | 180830 | 1920141 | 72.78 | 2.55 | 9.01 | 5.24 | 4.82 | 94.76 | 95.18 | 1.09 |
| FIN | 182757 | 198421<br>0 | 72.14 | 3.21 | 8.84 | 5.89 | 5.02 | 94.11 | 94.98 | 1.17 |
| GB<br>R | 195372 | 252572<br>1 | 72.63 | 2.88 | 7.53 | 5.21 | 3.79 | 94.79 | 96.21 | 1.37 |
| IBS | 181384 | 192422<br>7 | 72.64 | 2.65 | 9.04 | 5.07 | 4.73 | 94.93 | 95.27 | 1.07 |
| MX<br>L | 185618 | 173184<br>7 | 71.35 | 3.93 | 10.10 | 6.16 | 6.13 | 93.84 | 93.87 | 1.00 |
| PU<br>R | 184503 | 197439<br>0 | 72.17 | 3.35 | 8.94 | 2.85 | 3.14 | 97.15 | 96.86 | 0.91 |
| CL<br>M | 211683 | 259105<br>0 | 71.88 | 3.86 | 7.91 | 3.28 | 2.94 | 96.72 | 97.06 | 1.11 |
| PEL | 164146 | 151171<br>4 | 71.24 | 4.25 | 10.19 | 11.93 | 10.27 | 88.07 | 89.73 | 1.16 |
| GIH | 233021 | 208357<br>3 | 69.40 | 5.73 | 10.52 | 2.81 | 4.22 | 97.19 | 95.78 | 0.66 |
| PJL | 235204 | 193875<br>5 | 69.90 | 5.33 | 11.27 | 2.38 | 4.33 | 97.62 | 95.67 | 0.55 |
| BEB | 288816 | 260196<br>5 | 68.72 | 6.56 | 10.44 | 1.91 | 3.35 | 98.09 | 96.65 | 0.57 |
| STU | 242802 | 201763<br>7 | 68.84 | 6.24 | 11.22 | 2.38 | 4.41 | 97.62 | 95.59 | 0.54 |
| ITU | 243007 | 199722<br>8 | 68.37 | 6.16 | 11.33 | 2.66 | 4.51 | 97.34 | 95.49 | 0.59 |

We further looked at the individual- and population-level Neanderthal and Denisovan genome coverage as a function of sample size. The idea was to investigate how much of the Neanderthal or Denisovan genome could be recovered from a single or more individuals. Our hypothesis was that since the proportion of introgression is reported to be higher in East Asian individuals, then we should recover more archaic ancestry from a sample of East Asian individuals. To test this, we measured archaic genome coverage (see Methods) at various sample sizes to investigate the amount of Neanderthal variants that we could recover from a set of individuals. Figure 2A confirms that East Asians have more Neanderthal genome coverage per individual compared to individuals in other populations, consistent with previous studies [10,11,30,31]. For Denisovan variants, East Asian individuals exhibit similar levels of coverage as South Asian individuals (Figure 2B). When we look at the relationship between the amount of Neanderthal or Denisovan variants and sample size, we find that East Asians have nearly identical genome coverage to Europeans and admixed Americans as the sample size increases, and have lower coverage than South Asian populations (Figure 2F). Notably, in the case of Neanderthal-Unique variants, we actually recover more of the Neanderthal genome from a set of European genomes than a set of East Asian genomes, which is opposite of what we would expect from the findings at the individual level (Figure 2A-B). This suggests that while archaic variants in East Asians are found at higher frequency than in Europeans, more of these variants are shared between individuals in East Asia compared to Europe (Supp. Figure 7). For Denisovan-unique variants, we recover more from a set of East Asian individuals than Europeans which we expect given that Europeans exhibit almost no Denisovan ancestry. Perhaps most surprising is that we recover the largest amount of Neanderthal and Denisovan genome from any set of South Asian individuals even though South Asians have similar or lower individual-level genome coverage to East Asians (Figure C-D). The observation may point to a larger proportion of introgression from one or more introgression events from distinct archaic populations into the ancestral populations of South Asia or to more population structure in South Asian populations.

### Neanderthal ancestry in Europeans and East Asians

Figure 2B shows that at a sample size of 25 or larger, we recover more Neanderthal ancestry from Europeans than East Asians. If we compare the ratio of Neanderthal-Unique genome coverage between East Asians and Europeans, we observe an EAS/EUR ratio of 1.2 at the individual archaic genome coverage level, consistent with the 20% enrichment of Neanderthal ancestry reported in the literature [10,11,30,31]. However, as sample size increases, the EAS/EUR ratio approaches 1.01 (see Figure 4), and at the highest sample sizes, Europeans actually exhibit higher archaic genome coverage at the population level, with an EAS/EUR ratio of 0.97. This pattern is observed using archaic SNP data and we also recover the same signature using the introgression maps inferred for Europeans and East Asians using alternate methods [11,18,31] (Supp. Figure 4).

As several studies have suggested that East Asians have more Neanderthal ancestry due to more than one introgression event from Neanderthals, we wanted to assess whether one or two introgression events from Neanderthals into East Asians could lead to the observed pattern. Specifically, we simulated under a demographic model that accommodates up to two introgression events from Neanderthals into East Asian populations (see Methods and supplementary Figure 3). We varied two parameters representing differing proportions of one-pulse and two-pulse introgression models, ranging from a first pulse of 1% to 4% and a second pulse from 0 to 1% (Supp. Figure 3) for a total of 36 parameter combinations. We find that the parameters that minimize the mean squared error between the simulated and empirical EAS/EUR ratio curves correspond to a model with a first pulse of Neanderthal admixture of 3% and a second pulse of admixture into East Asians exclusively of 0.5% (Figure 4). Interestingly, several parameter combinations capture the observed pattern of the ratio being greater than 1 at n=1 and less than 1 at larger sample sizes, but none capture the exact shape of the empirical curve. The 5 best-fitting models have a second pulse that is 10-20% the magnitude of the first pulse, and only one of the 10 best-fitting models had only a single pulse of admixture. The worst-fitting models were any models with two pulses of admixture where the second pulse is ≥ 50% of the magnitude of the first (Supp. Table 1). Single-pulse models show a similar shape to the ratio curve observed in the empirical data, but the ratio in these models decreases more steeply with sample size, making for a poorer fit (Supplemental Table 1).

## Discussion

Our study of the apportionment of archaic alleles and of archaic genome coverage at the individual- and population-levels adds a new dimension to understanding the evolution of surviving archaic variation in modern human populations. We find that archaic variants in modern human populations are sufficient to recapitulate the population structure that is typically observed for East Asian, South Asian, European, and admixed American populations (Figure 3, Supp. Figure 2). Despite this regional grouping, there is more archaic variation that is shared between populations than population-unique (Table 2) — consistent with Lewontin’s [51] original observations, as well as more recent studies (e.g. [2, 51,52]). The only exception are Denisovan-unique variants, where the majority of alleles are unique to South Asia and to a smaller degree East Asia (Figure 3C). There is evidence of at least two distinct introgression events in the history of modern human populations, from highly diverged Denisovan-like populations, and Denisovan ancestry of early East Asians correlates with that in present-day East Asian and Austronesian populations, but not South Asian ones [18,20,54,55,63]. This suggest that it is possible that East Asian and South Asian populations received genomic contributions from distinct Denisovan populations. Interestingly, unlike other populations, archaic variation in South Asians tends to be at lower frequencies (Supplementary Figure 6), perhaps due to their complex history of mixtures between different ancestral groups that may have reduced the frequencies of archaic variants [53].

On the paradox of elevated Neanderthal ancestry in modern East Asians relative to Europeans, our results are consistent with previous findings [31,56,57], but only at the individual level. Remarkably, this difference in genome coverage is reversed at the population level. This suggests that the East Asian population has fewer Neanderthal introgressed segments than European populations but these segments are at higher frequencies, which results in higher Neanderthal genome coverage per individual. Conversely, the European population retains more Neanderthal segments, recovering a larger portion of the Neanderthal genome at the population level (Figure 2A, 2B). The retention of more unique Neanderthal variants in Europeans may certainly be related to modern demographic history, as East Asians experienced a more severe founder effect with a more rapid recovery [25,49,50]. For instance, we find that East Asian individuals tend to share archaic segments more often than Europeans as measured by homozygosity of tracts (Supplemental Figure 8). More natural selection acting on archaic variants in East Asians than in Europeans might also play a role in creating these patterns.

We used simulated datasets to test whether demographic hypotheses could explain how the ratio of Neanderthal genome coverage between East Asians and Europeans changes as a function of the sample size. In particular, we tested the number and magnitude of Neanderthal admixture events, while also taking inferred demographic differences between these two populations into account [49]. The parameter combinations that minimize the mean squared error correspond to a model with two pulses where the second pulse is approximately 10-20% of the magnitude of the first (see Figure 4), but these parameters fail to capture the full shape of the curve. Interestingly, both single and two pulse models can reproduce the feature of East Asians having more archaic coverage at an individual level, and Europeans having more coverage as the sample size increases, suggesting an attenuating effect of demography even in the case when the actual proportion of introgression is higher in East Asians. Models where the second pulse is at least 50% of the magnitude of the first pulse result in so much archaic genome coverage in East Asians that the ratio remains above one regardless of sample size, suggesting that increasing the proportion of introgression will not result in a better fit. Single-pulse models show a similar shape to the ratio curve observed in the empirical data, but the ratio in these models decreases more steeply with sample size, making for a slightly poorer fit (Supplemental Table 1).

While a model with two introgression events has the smallest mean squared error, none of our simple models perfectly reconstruct the EAS/EUR archaic coverage ratio curve (Figure 4), suggesting that more investigation of these demographic patterns is needed. We acknowledge that we have only considered a small number of parameter combinations, and further exploration of the parameter space may reveal combinations of first and second pulse proportions that provide an even better fit to the data. Additionally, there are demographic models we have not considered, such as an influx of unadmixed individuals into Europe from Northern Africa creating a “dilution” effect of archaic ancestry in modern Europeans [28], or the occurrence of Neanderthal admixture into Europeans as well as East Asians (a “three pulse” model). There is growing evidence of encounters between modern humans and various Neanderthal populations in geographically distinct regions of Eurasia (Fu et al. 2015; Zeberg et al. 2020; Taskent et al. 2020; Villanea et al. 2021; Hajdinjak et al. 2021). On the question of whether Europeans also received additional Neanderthal ancestry, recent evidence indicates the earliest Europeans encountered and admixed with distinct Neanderthal lineages but failed to leave descendants in today’s Europe (Oase-1 [59]), and some are more closely related to East Asian populations (Hajdinjak et al. 2021). These early Europeans were later replaced by human groups who only carried the original Neanderthal genomic ancestry shared by all Eurasians (Svensson et al. 2021).

Our study highlights how examining patterns of archaic variation in modern human variation can lead to insights on the evolution of archaic variation in humans. As a case in point, we find that our examination of South Asians reveals a rich and unique pattern of archaic ancestry. Previous studies comparing archaic ancestry in Eurasians have focused mostly on East Asians and Europeans [19,25,31,57], but our results suggest that South Asians have higher archaic genome coverage at the population level than both Europeans and East Asians (Figure 2). South Asians also display a large proportion of *rare* archaic alleles compared to other Eurasians (Table 1, Supplementary Figure 6), and a much larger number of unique archaic alleles compared to other populations (Table 2, Figure 4). Future inclusion of other South Asian and Oceanian populations may also help characterize the dynamics of Denisovan introgression, and modeling of archaic genome coverage accounting for periods of bottlenecks, expansions, gene flow and natural selection that followed the introgression events may reveal how evolutionary processes shaped the patterns of archaic ancestry in modern humans.

## Conclusions

By following in Lewontin’s steps and inspired by his classic 1972 study, we find new insights into modern population dynamics. Similar to Lewontin’s findings fifty years ago, we find that the largest component of archaic variants are shared in Eurasian populations, and the majority of archaic diversity is allocated to individual variation within populations. Summarizing archaic genome coverage at the individual- and population-levels allowed us to extract more information from the sharing and identity of archaic alleles, and use this information to test hypotheses of archaic admixture. Our results suggest that a model with a second Neanderthal introgression event into East Asians may explain observed differences in Neanderthal ancestry between East Asians and Europeans, suggesting that it is likely not solely due to differences in recent demographic history of these populations. Our analysis also shows that patterns of archaic variation in South Asian populations points to complex histories both of archaic introgression and more recent mixtures of ancestral groups that have shaped patterns of archaic variation differently than in Europeans or East Asians. Closer examination of how archaic genome coverage patterns change under a range of demographic models with the effects of natural selection will yield a better understanding of the population history of both modern and archaic humans.

## Acknowledgments

In memory of Richard Lewontin (1929-2021). EHS, FV and KEW were supported by NIH grant R35GM128946 (to Emilia Huerta-Sanchez). EL was supported by the James Watson Award from the Smurfit Institute of Genetics. The authors also thank Dr. Benjamin Peter for discussions and providing insightful comments related to this work.

## Author Contributions

KEW participated in study design, carried out the data analysis and simulations, and drafted the manuscript. FV participated in study design, assisted in creating the simulations and helped draft and revise the manuscript. EL participated in the data analysis and helped revise the manuscript. EHS conceived of the study, helped design the study, coordinated the study and helped draft and revise the manuscript. All authors gave final approval for publication.

