## Supplementary material for "On the Apportionment of Archaic Human Diversity": Supp. Materials

**Supplementary Figures and Tables**
**On the Apportionment of Archaic Human Diversity**

Kelsey E. Witt1,[2], Fernando Villanea[3], Elle Loughran[4], Emilia Huerta-Sanchez[1,2,4]

1 Ecology, Evolution, and Organismal Biology, Brown University
2 Center for Computational Molecular Biology, Brown University
3 Department of Anthropology, University of Colorado Boulder
4 Smurfit Institute of Genetics, Trinity College Dublin

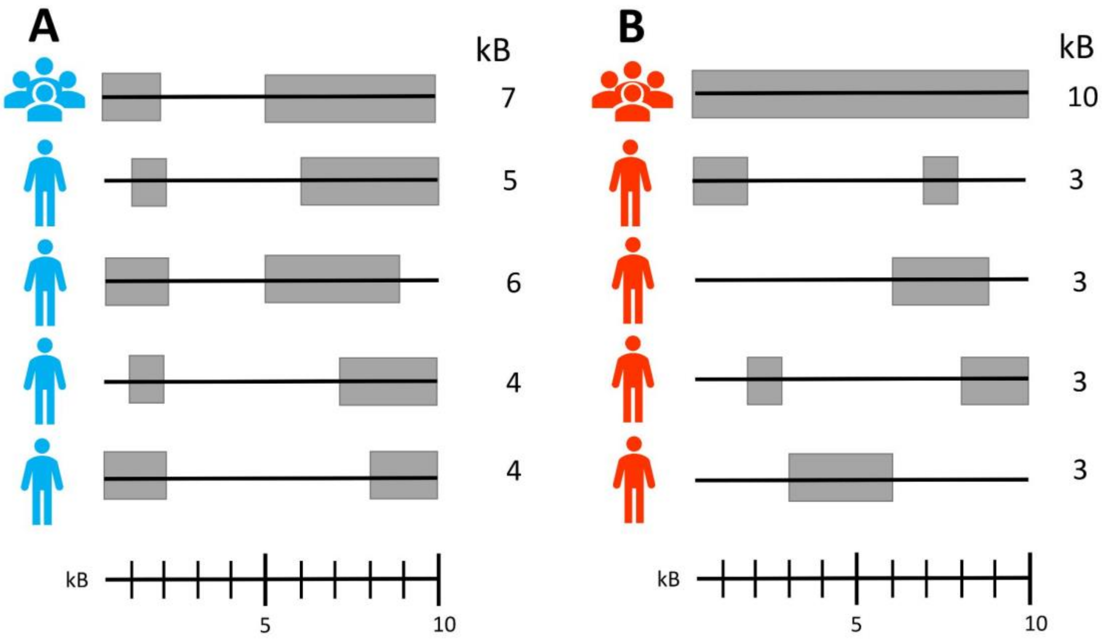


**Supplemental Figure 1:** An illustration of population- and individual-level genome coverage using archaic ancestry tract lengths. Here we show the ancestry tracts (gray boxes) present in a genomic region (the black line) for two populations, A and B. Each population contains four individuals, and their genome coverage is shown next to each individual along with the total amount of ancestry they have. For the population-level coverage, each archaic tract that is found in any individual in the population counts towards the total, so population-level coverage is the sum of archaic tracts found across all individuals in that population.

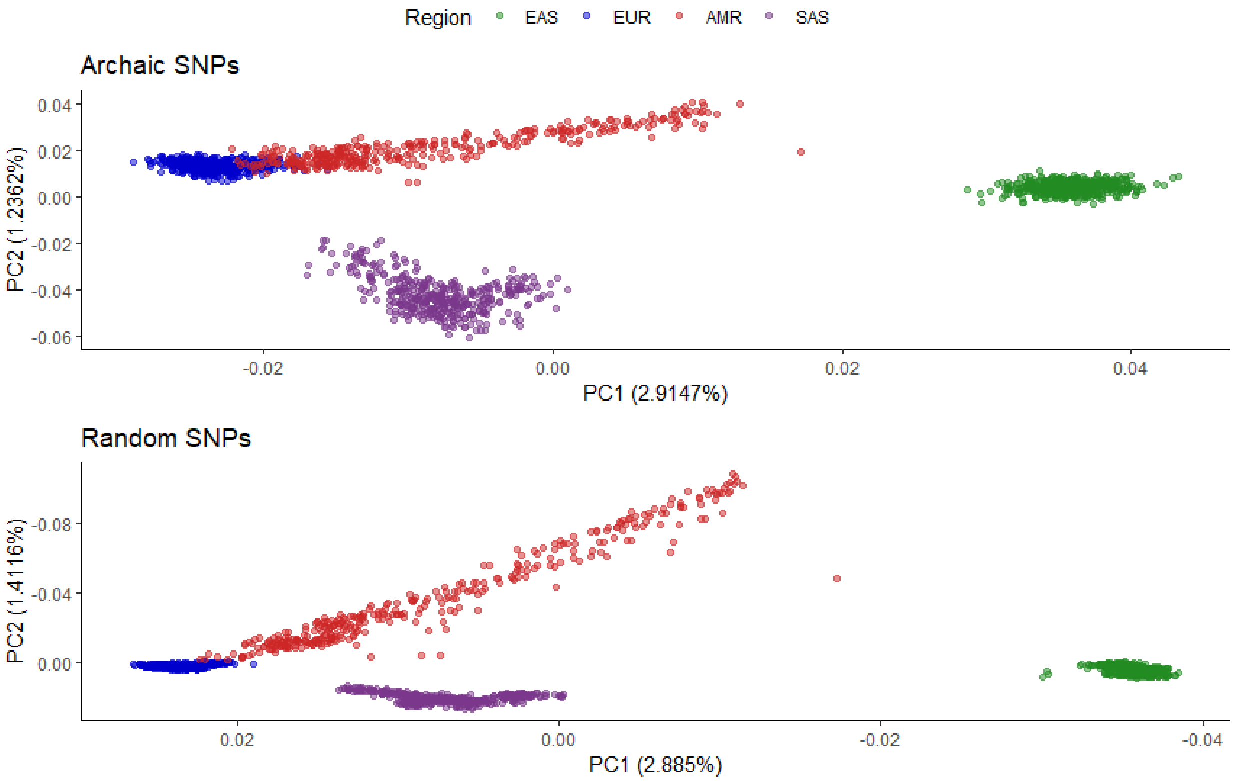


**Supplemental Figure 2:** A comparison of the PCA of the All-Archaic set compared to an equally-sized random sample of SNPs. In both cases, SNPs had to have a frequency of <1% in Africa but a frequency greater than at least one non-African population. The x and y axes on the random SNP plot have been flipped to allow for easier comparisons.

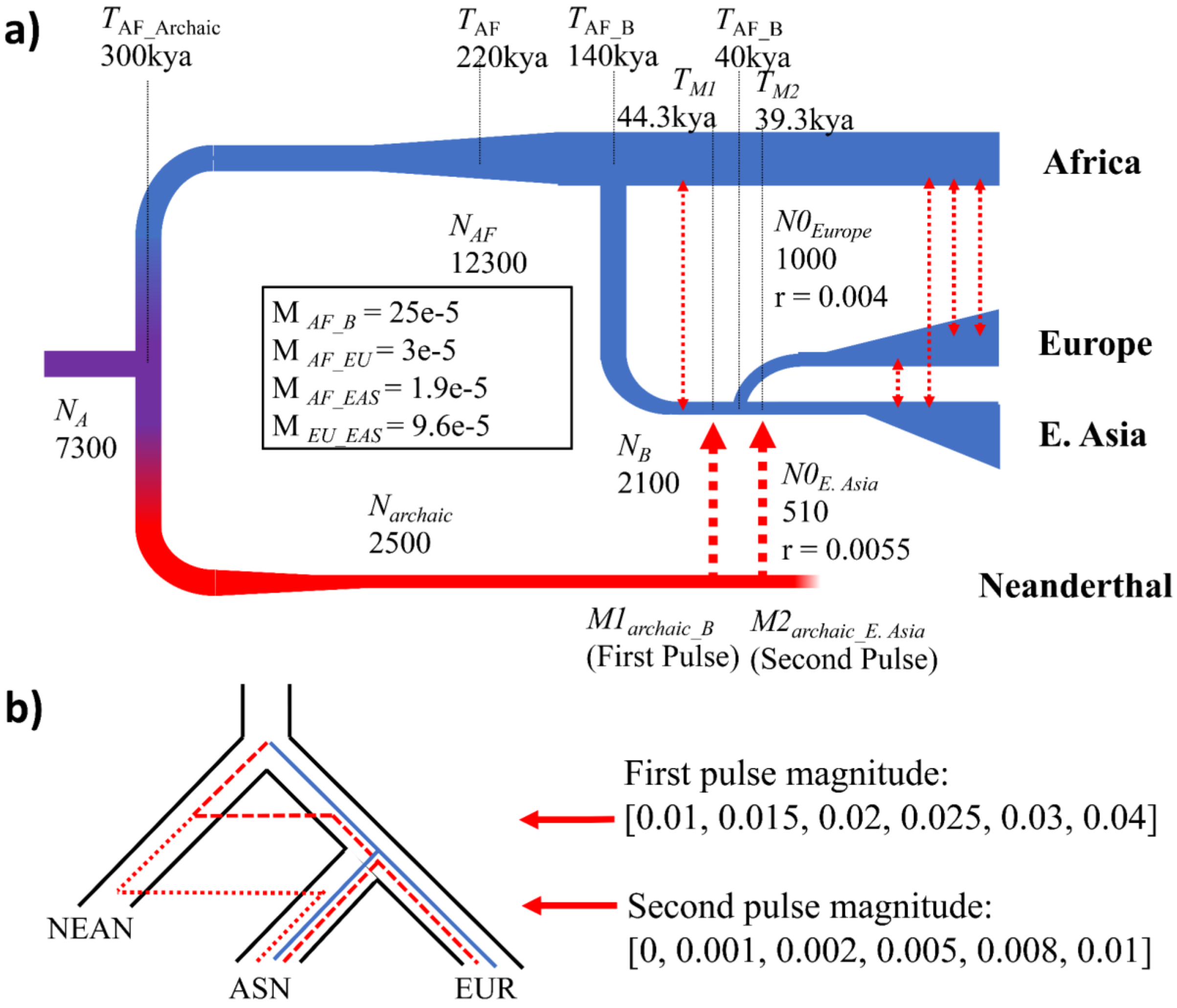


**Supplemental Figure 3**: The demographic model used in our simulations. a) A summary of the model, including population sizes and the timing of population splits and occurrences of gene flow. b) An illustration of the two pulses of gene flow, a “first pulse” from Neanderthals into the ancestor of East Asians (ASN) and Eurasians (EUR), and a “second pulse” from Neanderthals into East Asians. We used six values of gene flow for each pulse, which are specified in the brackets above.

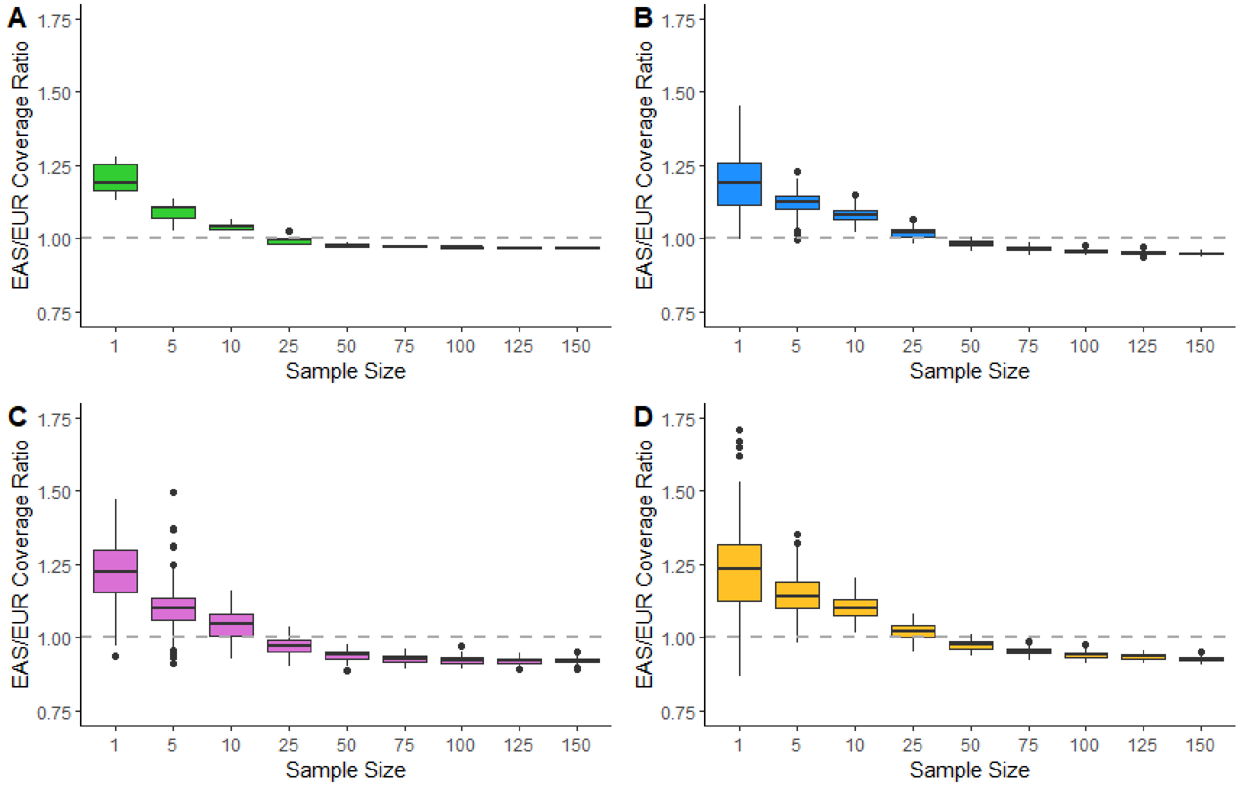


**Supplemental Figure 4:** A comparison of the ratio of genome coverage for East Asians/Europeans across varying sample sizes, using four different methods. A) SNPs with low frequency in Africa and shared with archaic populations, as calculated in this paper. B) SNPs identified using DICAL-ADMIX, as calculated by Steinrucken et al. 2018. C) SNPs identified using Sprime, as used in Browning et al. 2018. D) Ancestry tracts identified using a conditional random field method, as used in Sankararaman et al. 2014

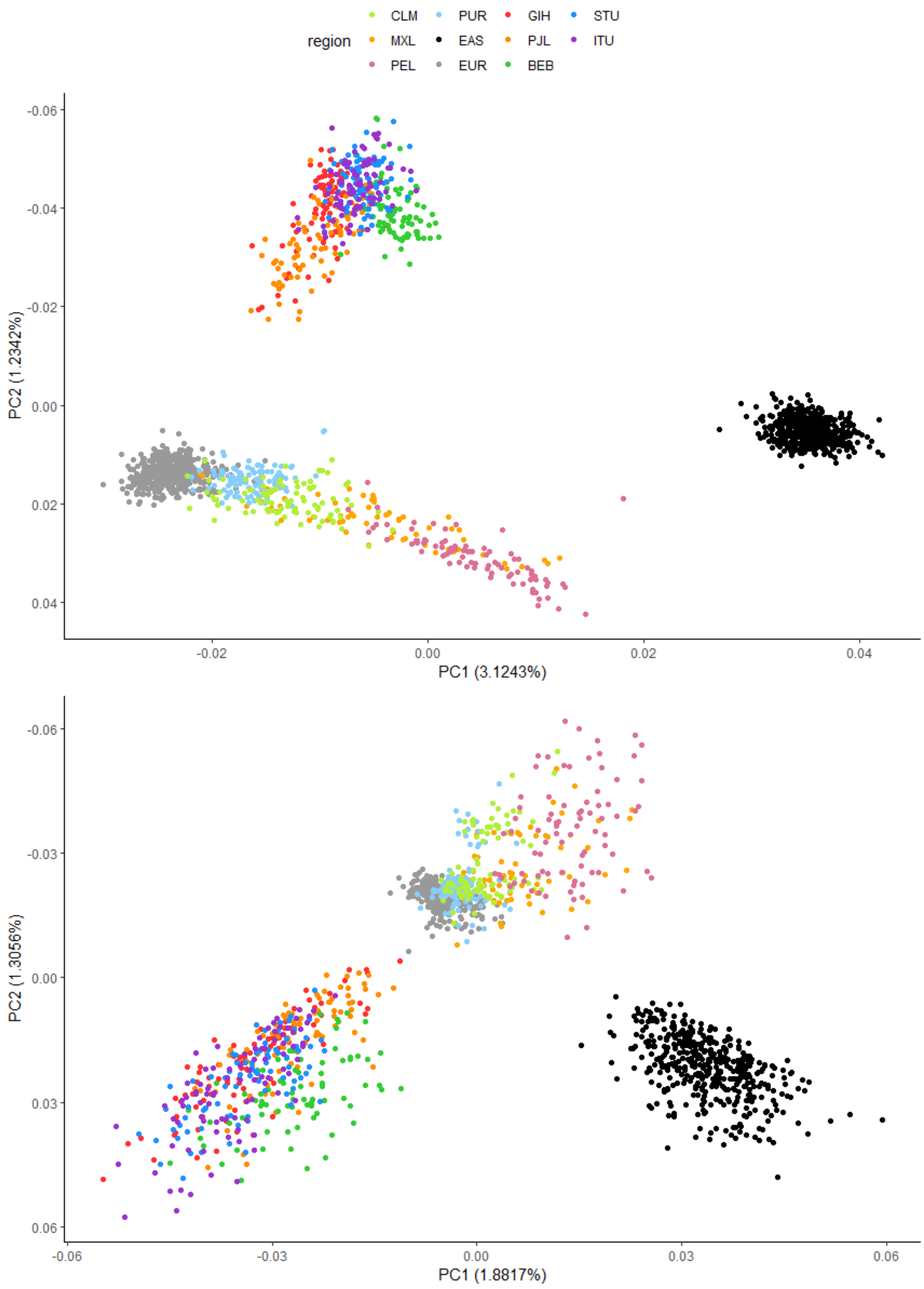


**Supplemental Figure 5:** A PCA of 1000 Genomes populations, using only Neanderthal-Unique (A) and Denisovan-Unique (B) SNPs, which are identical to Figures 2B-C but are re-colored to show the American and South Asian populations distinctly. East Asians (EAS) and Europeans (EUR) are color-coded by their super-population, while the rest are color-coded according to their specific population. Abbreviations follow standard 1000 Genomes conventions.

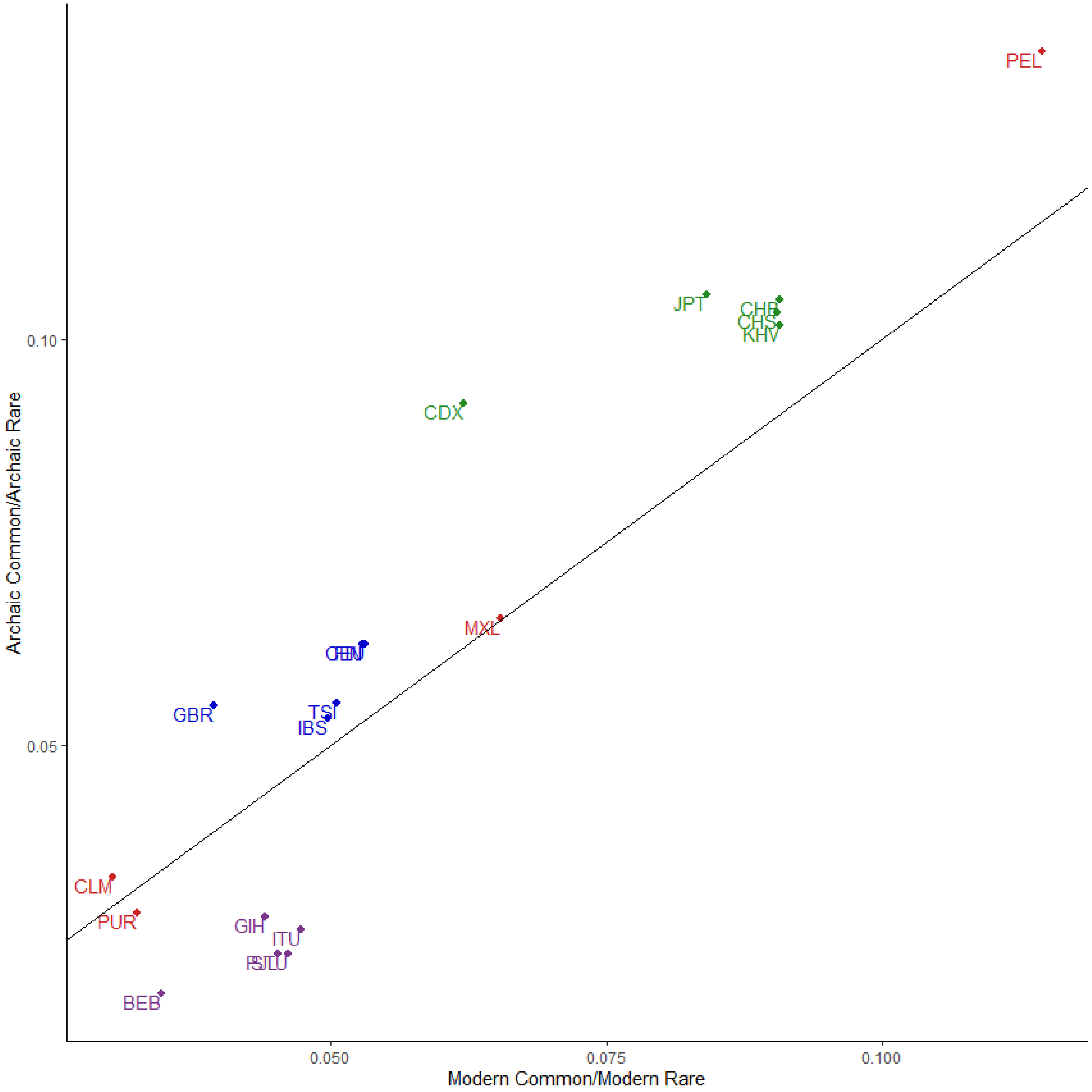


**Supplemental Figure 6:** Comparing the ratio of common (f>0.2):rare (0.1<f<0.2) alleles between "archaic" alleles (<1% frequency in Africa and shared with an archaic individual) and "modern" alleles (<1% frequency in Africa but not shared with an archaic individual). Each population is labeled according to standard conventions of the 1000 Genomes Consortium, and are color-coded by region (Americas - red, Europeans - blue, East Asians - green, South Asians - purple). The solid black line is the y=x line.

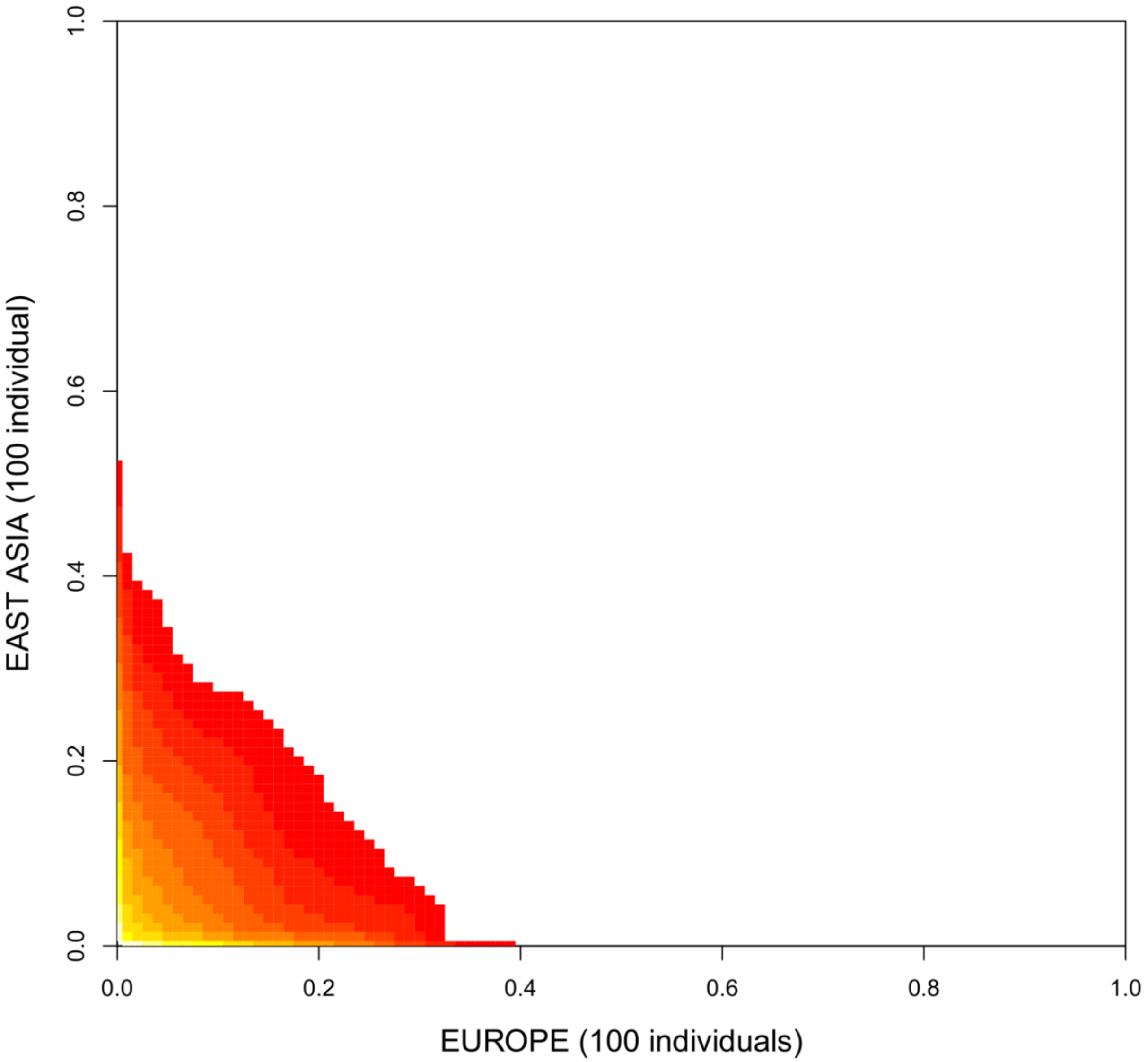


**Supplemental Figure 7:** Joint site-frequency spectrum for introgressed Neanderthals SNPs in the CEU (Europe) and CHS+CHB (East Asia) populations from the 1K Genomes Project, as called in Steinrucken et al. (2018). Each population was projected down to represent 100 individuals using a hypergeometric distribution.

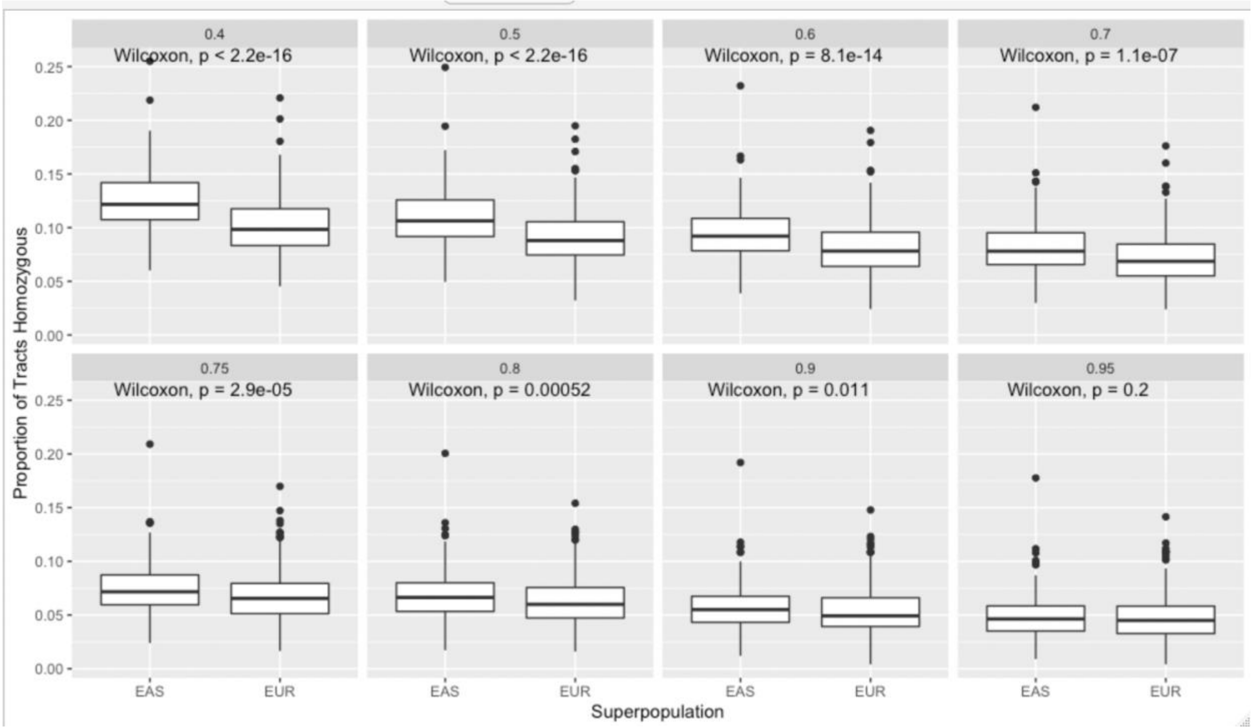


**Supplemental Figure 8**: A comparison of the proportion of homozygous archaic introgressed tracts between Europeans and East Asians, using the introgressed haplotypes identified in [11]. We used different threshold percentages of overlap between haplotypes to consider them as homozygous (40, 50, 60, 70, 75, 80, 90, 95%), and recorded the p-value for the difference between each population.

**Supplemental Table 1:** The mean genome coverage ratios for each demographic model and sample size, as well as the mean squared error relative to the empirical mean genome coverage. The models are listed in numerical order of mean squared error, from smallest (best fit) to largest (worst fit)

| First Pulse | Second Pulse | n=1 | n=5 | n=10 | n=25 | n=50 | n=75 | n=100 | n=125 | n=150 | mean squared error |
|---|---|---|---|---|---|---|---|---|---|---|---|
| 0.03 | 0.005 | 1.189696 | 1.08013 | 1.037286 | 0.996081 | 0.9825 | 0.978899 | 0.979075 | 0.980154 | 0.980708 | 0.001952 |
| 0.04 | 0.005 | 1.190141 | 1.08972 | 1.049135 | 1.010436 | 0.993524 | 0.989067 | 0.987945 | 0.988143 | 0.98854 | 0.001966 |
| 0.025 | 0.005 | 1.232237 | 1.117943 | 1.07307 | 1.028005 | 1.009236 | 1.002853 | 1.001505 | 1.000973 | 1.000538 | 0.002006 |
| 0.04 | 0.008 | 1.222972 | 1.12458 | 1.078652 | 1.032773 | 1.013989 | 1.00811 | 1.006295 | 1.005646 | 1.005291 | 0.00226 |

| | | | | | | | | | | | |
|---|---|---|---|---|---|---|---|---|---|---|---|
| 0.02 | 0.002 | 1.212881 | 1.06017 | 1.011283 | 0.969407 | 0.956572 | 0.956408 | 0.95788 | 0.959369 | 0.962114 | 0.002334 |
| 0.03 | 0.008 | 1.253686 | 1.141299 | 1.095401 | 1.047189 | 1.026402 | 1.01997 | 1.016884 | 1.015713 | 1.014729 | 0.003048 |
| 0.04 | 0.01 | 1.238628 | 1.143748 | 1.098182 | 1.052445 | 1.029149 | 1.021627 | 1.017948 | 1.015494 | 1.014841 | 0.003095 |
| 0.02 | 0 | 1.192763 | 1.043749 | 0.991961 | 0.946434 | 0.932039 | 0.932552 | 0.93532 | 0.937986 | 0.94072 | 0.003591 |
| 0.015 | 0.001 | 1.336263 | 1.064899 | 1.007043 | 0.950907 | 0.933982 | 0.932978 | 0.935231 | 0.938351 | 0.941319 | 0.003815 |
| 0.02 | 0.001 | 1.172018 | 1.035024 | 0.991952 | 0.952314 | 0.939576 | 0.940794 | 0.942624 | 0.945394 | 0.947957 | 0.003861 |
| 0.015 | 0.002 | 1.341031 | 1.161164 | 1.10759 | 1.04802 | 1.022349 | 1.014152 | 1.010943 | 1.008237 | 1.008018 | 0.003902 |
| 0.025 | 0.001 | 1.138964 | 1.027555 | 0.988079 | 0.950141 | 0.941187 | 0.941205 | 0.94419 | 0.94607 | 0.948357 | 0.004723 |
| 0.025 | 0.002 | 1.127792 | 1.026123 | 0.99282 | 0.956797 | 0.947858 | 0.949625 | 0.951114 | 0.953919 | 0.9554 | 0.004793 |
| 0.03 | 0.002 | 1.138864 | 1.023447 | 0.98094 | 0.943368 | 0.934587 | 0.936534 | 0.939607 | 0.943051 | 0.945944 | 0.005189 |
| 0.04 | 0.002 | 1.103585 | 1.015411 | 0.975194 | 0.94274 | 0.935982 | 0.937722 | 0.940901 | 0.944312 | 0.946744 | 0.006429 |
| 0.02 | 0.005 | 1.377634 | 1.19627 | 1.14004 | 1.074118 | 1.041206 | 1.030789 | 1.025635 | 1.022616 | 1.020832 | 0.006818 |
| 0.04 | 0.001 | 1.099022 | 1.00932 | 0.970541 | 0.933032 | 0.923787 | 0.924492 | 0.927704 | 0.931509 | 0.934214 | 0.007234 |
| 0.015 | 0 | 1.210976 | 1.00332 | 0.947733 | 0.904375 | 0.897684 | 0.901206 | 0.906955 | 0.911674 | 0.916616 | 0.007706 |
| 0.03 | 0.001 | 1.077713 | 0.98555 | 0.953095 | 0.921138 | 0.917665 | 0.921402 | 0.925734 | 0.930251 | 0.93343 | 0.009596 |
| 0.01 | 0 | 1.355756 | 0.996816 | 0.940513 | 0.895102 | 0.887004 | 0.889745 | 0.895517 | 0.900309 | 0.905462 | 0.010711 |
| 0.03 | 0 | 1.076084 | 0.978748 | 0.94067 | 0.909199 | 0.906695 | 0.912061 | 0.917286 | 0.921992 | 0.925797 | 0.010896 |
| 0.04 | 0 | 1.060928 | 0.9772 | 0.944925 | 0.917694 | 0.91574 | 0.920068 | 0.924794 | 0.928869 | 0.93251 | 0.010899 |

| | | | | | | | | | | | |
|---|---|---|---|---|---|---|---|---|---|---|---|
| 0.025 | 0.008 | 1.379493 | 1.235618 | 1.176334 | 1.110508 | 1.07407 | 1.059539 | 1.052094 | 1.047523 | 1.044433 | 0.011345 |
| 0.025 | 0 | 1.073125 | 0.96975 | 0.93334 | 0.905283 | 0.905387 | 0.910807 | 0.916476 | 0.921761 | 0.926161 | 0.011751 |
| 0.01 | 0.001 | 1.56168 | 1.148617 | 1.070835 | 1.003001 | 0.980194 | 0.976722 | 0.974872 | 0.975816 | 0.977218 | 0.012954 |
| 0.015 | 0.005 | 1.478102 | 1.250903 | 1.185793 | 1.114393 | 1.074212 | 1.059534 | 1.053072 | 1.047561 | 1.044742 | 0.016304 |
| 0.02 | 0.008 | 1.454097 | 1.276285 | 1.213061 | 1.135539 | 1.093366 | 1.074961 | 1.064605 | 1.058748 | 1.05442 | 0.018476 |
| 0.03 | 0.01 | 1.427209 | 1.291624 | 1.230863 | 1.152251 | 1.105619 | 1.085237 | 1.074164 | 1.067247 | 1.061685 | 0.019958 |
| 0.025 | 0.01 | 1.423983 | 1.28312 | 1.226276 | 1.156675 | 1.11464 | 1.095999 | 1.086386 | 1.079031 | 1.074142 | 0.021271 |
| 0.02 | 0.01 | 1.527358 | 1.364018 | 1.302263 | 1.224977 | 1.173259 | 1.14789 | 1.131678 | 1.121257 | 1.113336 | 0.044174 |
| 0.015 | 0.008 | 1.673963 | 1.42048 | 1.343313 | 1.245355 | 1.18294 | 1.154248 | 1.135966 | 1.124942 | 1.116816 | 0.063488 |
| 0.01 | 0.002 | 1.966776 | 1.348983 | 1.182326 | 1.100503 | 1.061109 | 1.046712 | 1.039144 | 1.035507 | 1.033566 | 0.071288 |
| 0.01 | 0.005 | 1.851656 | 1.408797 | 1.324064 | 1.225756 | 1.162817 | 1.13553 | 1.121965 | 1.111016 | 1.105094 | 0.078611 |
| 0.015 | 0.01 | 1.816151 | 1.538936 | 1.452146 | 1.339032 | 1.260354 | 1.222355 | 1.19864 | 1.182132 | 1.170338 | 0.117524 |
| 0.01 | 0.01 | 2.283249 | 1.764682 | 1.634202 | 1.476826 | 1.363079 | 1.305938 | 1.273433 | 1.250045 | 1.232124 | 0.283852 |
